# Population-specific recombination maps from segments of identity by descent

**DOI:** 10.1101/868091

**Authors:** Ying Zhou, Brian L. Browning, Sharon R. Browning

## Abstract

Recombination rates vary significantly across the genome, and estimates of recombination rates are needed for downstream analyses such as haplotype phasing and genotype imputation. Existing methods for recombination rate estimation are limited by insufficient amounts of informative genetic data or by high computational cost. We present a method for using segments of identity by descent to infer recombination rates. Our method can be applied to sequenced population cohorts to obtain high-resolution, population-specific recombination maps. We use our method to generate new recombination maps for European Americans and for African Americans from TOPMed sequence data from the Framingham Heart Study (1626 unrelated individuals) and the Jackson Heart Study (2046 unrelated individuals). We compare our maps to existing maps using the Pearson correlation between estimated recombination rates. In Europeans we use the deCODE map, which is based on a very large set of Icelandic family data (126,407 meioses), as a gold standard against which to compare other maps. Our European American map has higher accuracy at fine-scale resolution (1-10kb) than linkage disequilibrium maps from the HapMap and 1000 Genomes projects. Our African American map has much higher accuracy than an admixture-based map that is derived from a similar number individuals, and similar accuracy at fine scales (1-10kb) to an admixture-based map that is derived from 15 times as many individuals.

## INTRODUCTION

Recombination, or crossing-over, of chromosomes is essential for proper chromosome disjunction during meiosis. Recombination rates vary across the genome, tending to increase with decreasing chromosome length^1^, increase near the telomeres, particularly in males^2^, increase in regions with high GC content^2^, and increase in hotspots^3^, many of which are associated with the PRDM9 motif^4^. Recombination rates differ significantly between female and male meioses^1^, although sex-averaged maps are suitable for many analyses that involve historical recombination, including estimation of demographic history^5,6^, estimation of mutation rates^7,8^, estimation of haplotype phase^9–11^, genotype imputation^12,13^, and inference of local ancestry in admixed genomes^14–16^. Recombination rates also differ by age^2,17^ and by individual^17,18^.

There are four primary existing approaches to recombination rate estimation. The first is analysis of family data^2,17,19,20^. In order to estimate recombination rates at high resolution, extremely large numbers of meioses are required. One of the largest sources of such meioses is the deCODE Icelandic data^2^. Advantages of the family-based approach are that it can estimate sex-specific rates, and that it allows investigation of individual-specific factors influencing recombination rates^2,17^. A disadvantage is that the large family databases required by this approach are rare, so population-specific rates are not available for most populations.

A second approach is sperm-typing, with recombination events identified by comparing haplotypes between sperm cells obtained from the same individual. This approach can be used to locate recombination hotspots^21,22^ and construct individual genome-wide recombination maps in males^23^. However, this approach has not been used to construct population-level recombination maps because it is not applicable to females, and because large databases of whole-genome sperm sequence are not available.

A third approach uses admixed genomes such as those from African Americans^24,25^. The local ancestry (i.e., continental origin of the genetic material at each point in the genome) is inferred, and positions of change in local ancestry are positions at which post-admixture recombination has occurred. This approach can use data from unrelated individuals, and each individual provides information from many meioses. One limitation of this approach is that it is only applicable to admixed populations. Another disadvantage is that it relies on local ancestry calls which can be inaccurate in some cases^26,27^.

A final approach uses linkage disequilibrium (LD) between loci^28–32^. Correlations between nearby alleles are broken down over many generations due to recombination, and thus there is a close relationship between LD and recombination rate. An advantage of LD-based estimation is that it is based on very large numbers of meioses, reaching far back into the past. However, if recombination rates have changed over time, the estimates will be averages across time rather than reflecting current rates, which may be a disadvantage for applications such as local ancestry inference that are based on recombination in recent generations. LD-based estimation is computationally challenging, and is also biased if an incorrect demographic history is assumed^31,32^.

We present a new approach based on estimated segments of identity by descent (IBD) in population samples. IBD segment ends represent points at which past recombination has occurred. Since the IBD segments result from shared ancestry in the past several hundred generations, the estimates of recombination rates reflect recent rates while incorporating information from a large number of meioses. Our approach is computationally efficient so that it can be applied to samples of thousands of individuals, resulting in highly precise estimates. When applied to samples from distinct populations, our approach provides population-specific rates of recombination.

## RESULTS

### Method overview

IBD segment endpoints are positions of past recombination events. The density of endpoints of IBD segments originating from common ancestors more recent than a reference time point is thus proportional to the recombination rate. We use this relationship to estimate relative recombination rates based on the endpoints of IBD segments.

There are two main challenges that must be addressed: inaccurate estimation of IBD endpoints and the unknown time to the most recent common ancestor. Because of genotype error and phasing error, IBD segment endpoints can be incorrectly determined. In our method, we apply a gap-filling strategy to address inaccurate IBD endpoints. When two or more IBD segments from the same pair of individuals are separated by only a small gap, and the gap contains very few (at most one for the analyses presented here) discordant homozygous genotypes, we merge the segments into a single segment^33^. This strategy is very efficient at removing incorrect IBD endpoints, even in the presence of significant genotype error (see simulation results below).

In addition, when we detect an IBD segment, we don’t generally know the number of generations to the most recent common ancestor, and in some regions we may detect IBD segments due to more distant ancestry, leading to higher rates of detected IBD segments in those regions. If the genetic lengths of the segments were known, we could filter the IBD segments by genetic length, as a proxy for age, and thus obtain uniform rates of detected segments across the genome. When the true genetic map is unknown, the genetic lengths of the segments cannot be used as a filtering criteria, and we use physical lengths instead. However, the distribution of IBD segments of a given physical length varies greatly. Regions with lower recombination rate will have more segments exceeding a physical length threshold than regions with higher recombination rate. If this difference is not considered, recombination rate will be overestimated in low-recombination regions and underestimated in high-recombination regions.

In order to address the issue of uneven IBD coverage, we use an iterative approach. Given the current estimate of recombination rates across the chromosome (the initial estimate has a constant rate of recombination), we obtain estimated genetic lengths of all IBD segments. We then selectively remove shorter segments in regions with higher rates of IBD segments until IBD coverage across the chromosome is approximately equal. To achieve this coverage equalization, we first divide the chromosome into intervals of equal physical length, and place within each interval the IBD segments that cover all or part of the interval. We determine the smallest number of IBD segments in any interval, and we remove the shortest (in estimated genetic length) IBD segments from each interval to reduce the number of segments in the interval to that smallest number. After this procedure, each interval contains the same number of IBD segments (Figure 1). After the coverage equalization, we count the remaining IBD endpoints within each interval to estimate the relative recombination rate for the interval. We repeat the procedure using the updated estimates of recombination rate. We find that 20 iterations suffices for accurate estimation.

**Figure 1:**
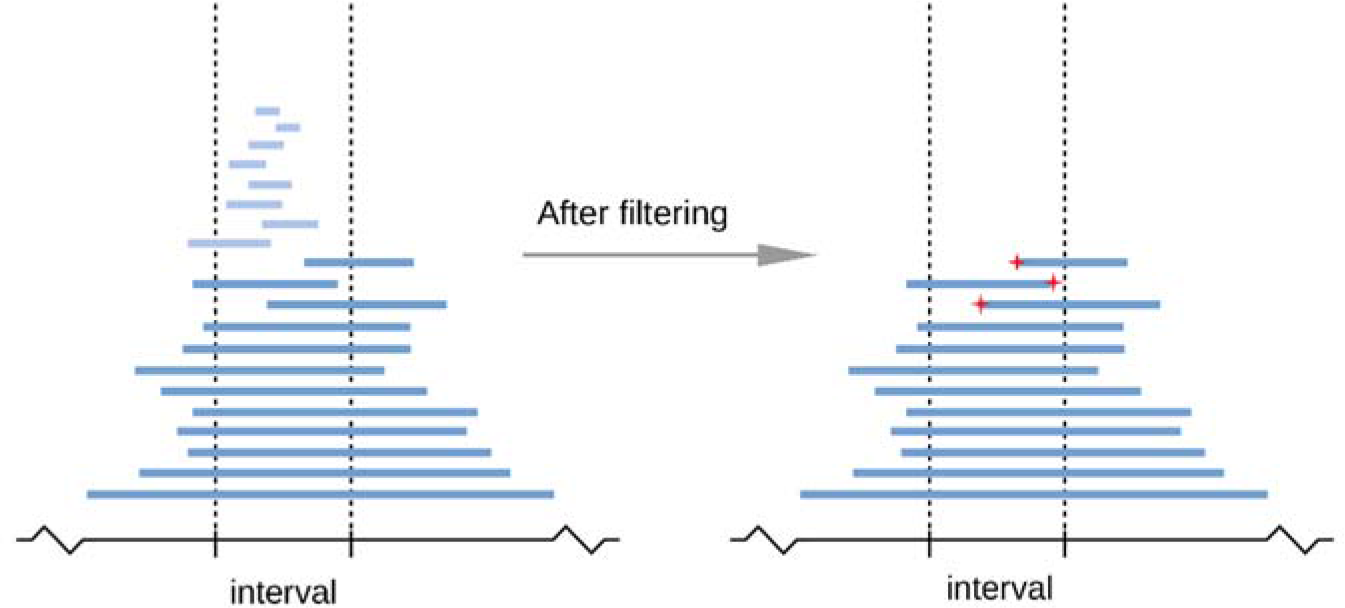
An illustration of the procedure for enumerating IBD endpoints for recombination rate estimation. In each iteration, IBD segments with short estimated genetic length are filtered out to achieve the required level of IBD coverage in the target interval, which is delineated with vertical dashed lines. In this example, segments in light blue are filtered out and the three remaining IBD endpoints falling within the interval (marked by red stars) are counted as recombination events corresponding to this interval.

### Validation by simulations

To evaluate whether our method produces unbiased estimates of recombination rate, we simulated data using a podium-like recombination map (Figure S1) and added genotype error (Methods). When the genotype error rate is low (0.01-0.1%), the average across 100 replicates of the estimated recombination rate matches the true recombination rate; when the genotype error rate is 0.5%, some bias is observed. With high quality sequence data, the genotype error rate for SNPs passing quality control filters is around 0.02%^34^. Our estimates are slightly inflated at the chromosome ends. This is because genotype error near the chromosome ends result in partial IBD segments that are too short to be detected and merged with other partial IBD segments by our gap-filling process. In our tests, the region with inflated recombination rates is generally shorter than 1Mb when the genotype error is ≤0.1%. We recommend that normalization of relative recombination rates using an external map be calculated using the central portion of the chromosome, excluding 1 Mb on each end.

We also used simulation to evaluate the precision of our method, first assessing the impact of sample size and resolution. The resolution, which we refer to as “scale” is the size of the intervals in which recombination rate is estimated. For example, with a 10 kb scale, the recombination rate is estimated in intervals of size 10 kb, and the resulting map has genetic positions at grid points that are 10 kb apart. We simulated 250 individuals, 500 individuals, and 1000 individuals under a Wright-Fisher model with constant effective population size (Ne = 10000). The recombination map used for this simulation is the Hapmap II combined LD map on chromosome 1:10 Mb-110 Mb.^35^ Pearson correlation coefficients between the estimated rates and the true rates across intervals increase for larger interval sizes and larger sample sizes (Figure 2). For the largest sample size, we obtain correlation coefficients over 0.9 for resolutions of 10 kb or greater and genotype error rates ≤0.1%. With smaller sample sizes, we obtain correlation coefficients over 0.9 for resolutions of 50 kb or greater and genotype error rates ≤0.1%.

**Figure 2:**
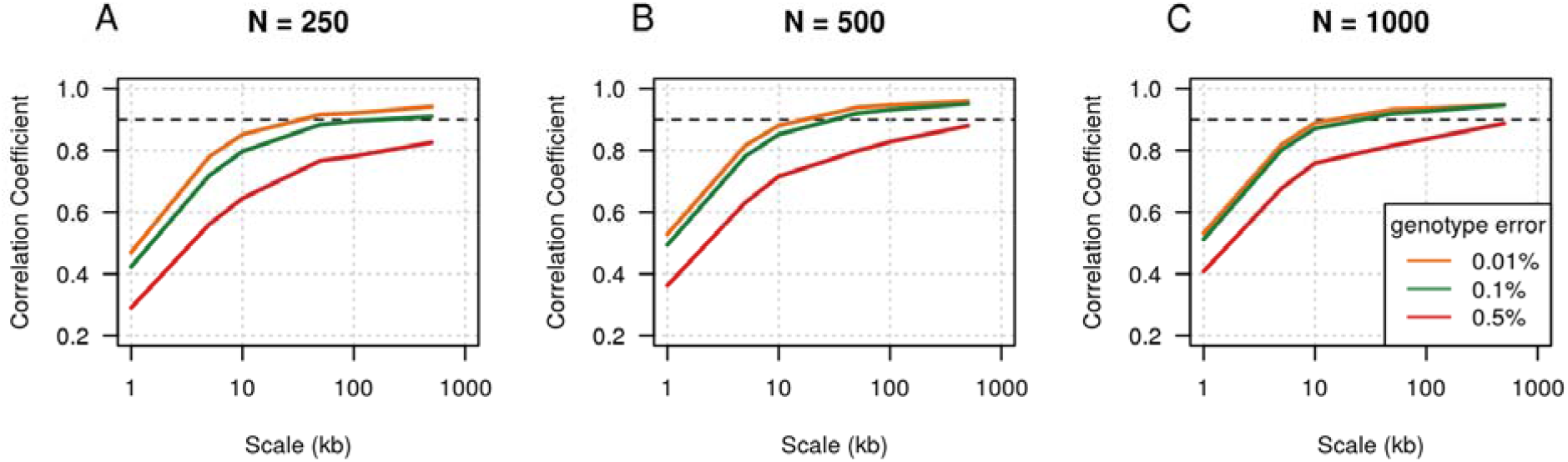
Pearson correlation coefficients between estimated recombination rates and true rates. 100 Mb of data were simulated using the Hapmap II combined LD map and a constant effective population size of 10,000. Sample sizes were 250 (panel A), 500 (panel B) or 1000 (panel C). Simulations with different genotype error rates have different colors. The x-axis gives the estimation scale (size of intervals in which recombination rates are estimated and for which correlation coefficients are calculated) on a log scale. The black dashed line shows a correlation coefficient of 0.9 for reference.

### Comparison with admixture-based recombination rate estimation

We simulated genotype data from an admixed African American demographic model^33^ in order to compare our IBD-based approach with RASPberry^25^, an admixture-based approach. We simulated 2500 admixed individuals and 100 individuals from each reference population (representing European ancestry and African ancestry). RASPberry uses the reference individuals to call local ancestry in the admixed individuals. Because RASPberry is computationally intensive, we simulated a 20 Mb region rather than 100 Mb. The recombination map in the simulation is the HapMap II combined LD map, chr1:10 Mb-30 Mb.^35^ We added genotype errors to the admixed and reference individuals and phased the data with Beagle 5.0 (Methods) before running the analyses. Our IBD-based method was applied to the admixed data only.

RASPberry uses the HapMix algorithm for ancestry inference, which analyzes each admixed individual independently and allows for parallelized computation over admixed individuals^15,25^. To reduce RASPberry’s wallclock compute time, we divided the 2500 admixed individuals into 250 sets of 10 individuals. We analyzed the data using a compute server with two 6-core Intel Xeon E5-2630 2.6 GHz processors and 128 GB memory running CentOS Linux. RASPberry required 20.1 cpu hours on average to estimate the ancestry switches for each set of 10 individuals. For comparison, our method required 11.1 cpu hours (1.0 hour of wall clock time, multi-threaded) to call the IBD segments, fill IBD gaps, and estimate the recombination map for the whole set of 2500 admixed individuals.

In assessing the accuracy of the estimates, we trimmed 5 Mb from each end of the simulated region (Figure 3) before computing the Pearson correlation coefficient with the true recombination rates, because accuracy is reduced near chromosome ends (results without this trim are shown in Figure S2). Estimates from our IBD-based method (IBDRecomb) have higher correlation with the true recombination rates than RASPberry at all scales. At a 10kb scale, our method’s estimates have a correlation coefficient around 0.9, while RASPberry’s correlation coefficient is around 0.65 (Figure 3). A likely explanation for the higher accuracy of our method is that our method utilizes recombination events before and after admixture, while RASPberry can only use recombination events that occurred after admixture.

**Figure 3:**
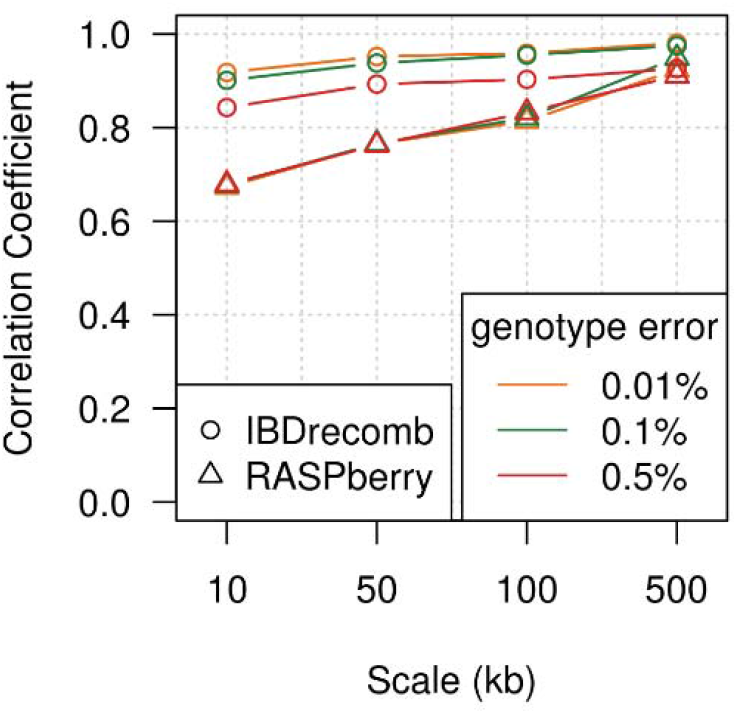
Pearson Correlation coefficients between estimated recombination rates and true rates. The results are based on simulations with different levels of added genotype error. Each end of the region was trimmed by 5 Mb before calculating correlation coefficients.

### Constructing a fine scale recombination map for the Framingham Heart Study data

We refer to the recombination rates that we estimated from the TOPMed Framingham Heart Study data as the FHS map. We compare the FHS map to other existing recombination maps, including the deCODE map based on Icelandic pedigrees^2^, the LD-based combined map from Hapmap II^35^, and LD-based maps from the 1000 Genomes Project^36^. Since most of these existing maps are available in the Genome Reference Consortium Human Build 37 (GRCh 37), we lifted over the deCODE map from build 38 to build 37, removing intervals not conserved between two genome builds (Methods).

Examination of a region on chromosome 1 shows that our FHS map captures the same hotspots that are found with other methods (Figure 4). For a genome-wide comparison, we calculate correlation coefficients between maps. We regard the deCODE map as the “gold standard” in our comparison of recombination maps estimated from Europeans, because this European-specific map is based on directly observed recombination events from a very large number of meioses. We calculate the Pearson correlation coefficient between a map’s recombination rate estimates and the deCODE map’s rates. In order to calculate the correlation coefficient at a given scale (such as 1kb), we divide the genome into intervals of this length, and obtain the estimated recombination rate for each such interval for every compared map. Since each map covers a slightly different subset of the genome, we ignore intervals that are not fully covered by all maps included within a given comparison. We excluded Bhérer et al’s refined maps^19^ from the comparison because Bhérer et al used the deCODE recombination events in their estimation. We find that all the European-ancestry maps have similar correlation coefficients to the deCODE map at scales ranging from 50 kb to 500 kb, while our FHS map has the highest correlation coefficients at scales ranging from 1kb to 10kb (Figure 5). Our map has higher correlation coefficients than the European LD-based maps at fine scales, indicating that our method can provide superior recombination rate estimation over LD-based methods.

**Figure 4:**
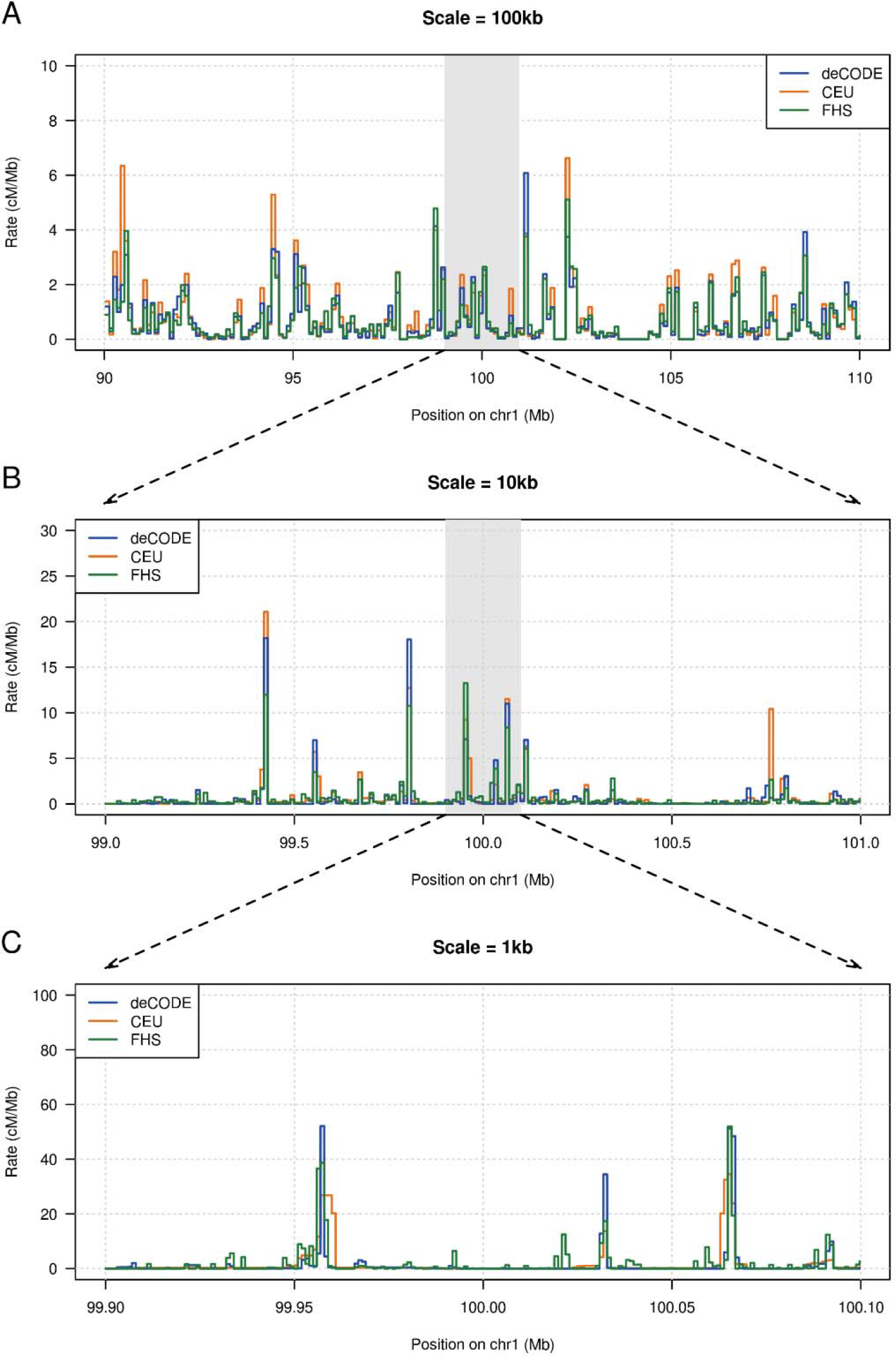
Estimated European recombination rates around chr1:100Mb. A) 20Mb at 100kb scale; B) 2Mb at 10kb scale; C) 200kb at 1kb scale. The three maps represent three different methods: pedigree-based (deCODE), LD-based (CEU from the 1000 Genomes Project), and IBD-based (FHS).

**Figure 5:**
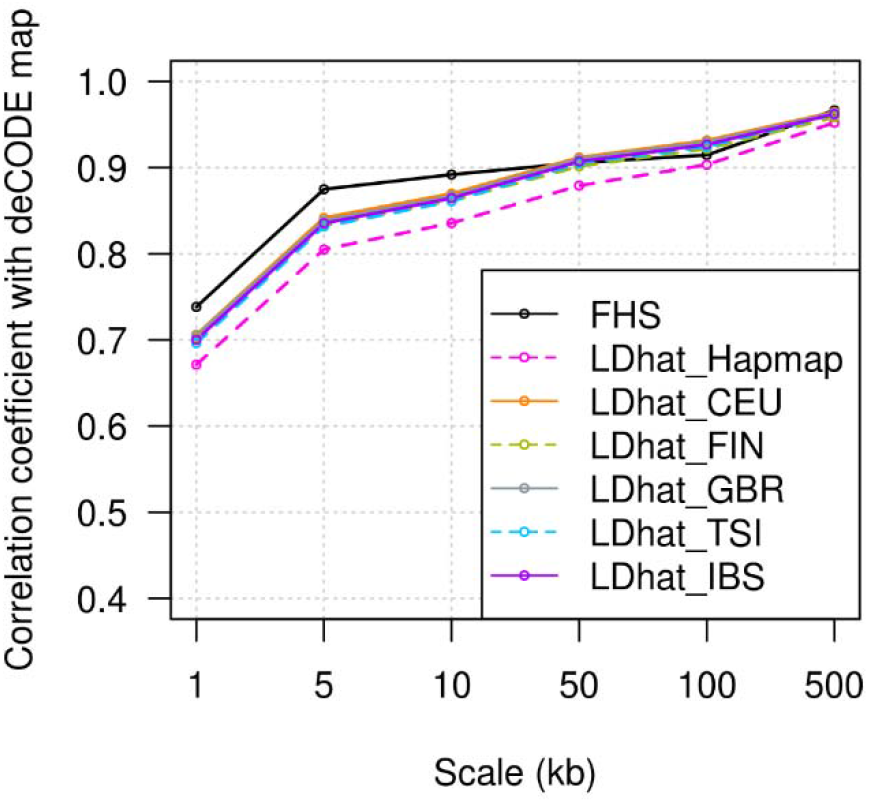
Pearson correlation coefficients between each map and the deCODE map at different scales. FHS is our IBD-based map from the TopMED Framingham Heart Study data. LDhat_Hapmap is the LD-based Hapmap combined map, while LDhat_CEU/FIN/GBR/TSI/IBS are the LD-based 1000 Genomes maps for the Utah residents with Northern and Western European ancestry (CEU), Finnish in Finland (FIN), British in England and Scotland (GBR), Toscani in Italia (TSI), and Iberian in Spain (IBS) populations, respectively.

### Constructing a fine scale recombination map for the Jackson Heart Study data

We also used our method to construct a recombination map for African Americans based on the data from the TOPMed Jackson Heart Study data. We compare our map (the JHS map) with three other maps constructed with African American data: the AA map^24^, the AfAdm map^25^, and the 1000 Genomes LD-based map for the ASW (Americans of African ancestry in SW USA) population^36^. The AA and AfAdm maps are constructed using counts of ancestry switches in 30,000 and 2864 admixed African Americans, respectively. We also compare to a 20%:80% weighted average of the 1000 Genomes LD-based maps for CEU (Utah residents with northern and western European ancestry) and YRI (Yoruba in Ibadan, Nigeria).

Again examining a region on chromosome 1, we find that our JHS map includes the same recombination hotspots found by other LD-based and admixture-based methods (Figure S3). For a genome-wide comparison, we calculate correlation coefficients between maps. The AA map, the JHS map, the LD-based ASW map, and the weighted CEU+YRI map are highly correlated at large scales (Pearson correlation coefficients > 0.85 at scales ≥50kb), and slightly different at fine scales (Table 1, Table S1). At scales ranging from 1kb to 10kb, the CEU+YRI map has the best performance with highest correlation coefficients to each other map (Table S1). It is even better than the ASW map which is inferred from an African American population, possibly because populations in the Americas experienced founding bottlenecks^33^ which reduce the number of unique historical recombination events represented in current-day genomes from these populations.

**Table 1:** Pearson correlation coefficients between estimated recombination rates for five African American genetic maps at different scales. The CEU+YRI map is the 20%:80% weighted average of the 1000 Genomes Project CEU and YRI maps. Results at other scales from 1kb to 500kb can be found in Table S1.

| Scale |  | AA | JHS | AfAdm | ASW | CEU+YRI |
| --- | --- | --- | --- | --- | --- | --- |
| 5kb | AA | 1.00 | 0.79 | 0.36 | 0.78 | 0.81 |
|  | JHS | 0.79 | 1.00 | 0.36 | 0.79 | 0.83 |
|  | AfAdm | 0.36 | 0.36 | 1.00 | 0.39 | 0.38 |
|  | ASW | 0.78 | 0.79 | 0.39 | 1.00 | 0.91 |
|  | CEU+YRI | 0.81 | 0.83 | 0.38 | 0.91 | 1.00 |
| 100kb | AA | 1.00 | 0.90 | 0.81 | 0.91 | 0.93 |
|  | JHS | 0.90 | 1.00 | 0.79 | 0.87 | 0.89 |
|  | AfAdm | 0.81 | 0.79 | 1.00 | 0.81 | 0.82 |
|  | ASW | 0.91 | 0.87 | 0.81 | 1.00 | 0.96 |
|  | CEU+YRI | 0.93 | 0.89 | 0.82 | 0.96 | 1.00 |

At fine scales (1-10kb), the JHS map and the admixture-based AA map have similar correlation with other maps, while at large scales (50-500kb) the AA map has higher correlation (Table 1, Table S1). Since the AA map is based on SNP array data, it is not surprising that it has lower relative accuracy at fine scales, while its large sample size (around 15 times as many individuals as in our JHS analysis) gives it high accuracy at large scales. Both the JHS map and the AA map have much higher correlations than the admixture-based AfAdm map to other maps at all scales. The AfAdm map is based on data with a sample size that is similar to that of our JHS data (2864 individuals for the AfAdm map and 2046 individuals for our JHS map). Hence it is notable that our JHS map has much better accuracy than the AfAdm map.

## DISCUSSION

We have presented the first IBD-based recombination rate estimation method, along with estimates of recombination rates in European Americans and African Americans. Our approach and maps have significant advantages over existing approaches and maps. Our approach is applicable to large population-based samples with sequence data, enabling the generation of high-resolution population-specific recombination maps. Our maps constructed from the Framingham Heart Study and the Jackson Heart Study will be useful for downstream analyses that require recombination maps, including haplotype phase estimation, genotype imputation, inference of demographic history, and inference of local ancestry in admixed individuals.

As with other indirect methods (admixture-based or LD-based estimation), our method requires the total genetic length of chromosomes from direct (family-based) estimation in order to convert relative recombination rates to absolute recombination rates. While family-based estimation of high-resolution genetic maps requires very large numbers of informative meioses, obtaining the approximate genetic length of a chromosome requires many fewer meioses. In addition, while recombination rates may change at small scales due to changes in hotspots, large-scale rates are conserved across populations^37^. Thus chromosome lengths from the Icelandic deCODE map (for example) may be used to normalize IBD-based relative recombination rates estimated in other human populations.

Generation of new maps with our method is straightforward, and we provide software to do so (Online Resources). Our method is applicable to humans and to other diploid species. With reductions in sequencing costs, it is likely that there will soon be suitable data for a variety of species, including model organisms, domesticated species, and wild species. The generation of high-resolution maps will facilitate other analyses in these populations. As input, our method requires high-quality genotype data (variant calls) on at least several hundred individuals, and a high-quality genome build for determination of physical positions. Sequence data is needed for accurate fine-scale estimation, but array data is adequate for estimation at large scales (Figure S4).

Our IBD-based method gives greater resolution than ancestry-switch based methods for constructing recombination maps from admixed individuals. This is because our method can detect recombination events that occurred before admixture, as well as those that occurred after admixture, while ancestry-switch based methods only use recombination between different ancestry segments that occurred after admixture. We built an African American recombination map with 2,046 unrelated African American individuals from the TopMED Jackson Heart Study data, which had significantly better accuracy than the admixture-based map constructed on 2,864 unrelated African American and Afro-Caribbean individuals, and similar accuracy at fine scales to an ancestry-switch based map constructed from 15 times as many samples (n = 30,000).

We also built a recombination map using data from the TopMED Framingham Heart Study data (n=1626), which represents a European-American population. This map shows better accuracy at fine scales (1-10 kb) than the LD-based maps for the 1000 Genomes Project European populations. Like our method, LD-based methods are based on past recombination events, however our method depends on recent recombination events, while LD-based methods are primarily based on recombination events occurring in the much more distance past. In contrast, family-based methods use recombination events from the past few generations. Recombination rates evolve over time^38^, so restricting the analysis to more recent events is advantageous for some applications.

Current recombination rates in Europeans and other out-of-Africa populations may differ from rates in African populations because of drift that occurred in the out-of-Africa bottleneck. For example, non-African populations predominantly carry the A allele of PRMD9, while African populations carry that allele at a frequency of only around 50%^4^. Thus LD-based recombination rates, which are based on older recombination events, may be more appropriate for African populations than for out-of-Africa populations. Indeed, while the LD-based maps for the 1000 Genomes European populations had inferior accuracy to our IBD-based map, LD-based maps from the 1000 Genomes African populations (the ASW map and the weighted CEU+YRI map) provided slightly superior accuracy to our IBD-based map. With larger sample sizes, we anticipate that our IBD-based approach could provide better maps than the LD approach, which is limited by computational cost to relatively small sample sizes.

The IBD-based approach has some limitations. The major obstacle to achieving higher accuracy at fine scales for our method is the difficulty in accurately establishing the exact IBD endpoints. Wrongly placed IBD endpoints may lead to false recombination rate peaks at fine scales and may also lead to underestimation in recombination hotspots. Currently, IBD estimation methods do not provide a representation of the uncertainty around the exact IBD endpoints. Future work could address this issue.

## METHODS

### Data processing

We used a coalescent-based simulator, msprime^39^, to simulate genetic data under different scenarios. We removed the phase information from the simulated haplotypes and added genotype error. Given a genotype error rate ϵ, and considering each genotype in turn, we added an error to the genotype with probability ϵ. When adding an error to a genotype, we selected one of the genotype’s two alleles at random, and changed that allele to its alternative form (all simulated markers are biallelic). Then we filtered sites to keep those with minor allele frequency larger than 0.05, and phased the data with Beagle 5.0^9,13^ (version 04Jun18.a80).

We applied our method to TOPMed whole genome sequence data from the Framingham Heart Study (FHS, download from dbGaP, phs000974.v2.p2), and the Jackson Heart Study (JHS, downloaded from dbGaP, phs000964.v2.p1). The individuals in the FHS data are European Americans, while the individuals in the JHS data are African Americans. To control genotype error, we only used biallelic SNPs passing all quality filters and with minor allele frequency larger than 0.05. We used Beagle 5.0^9,13^ (version 04Jun18.a80) to infer haplotype phase for each data set. We then used King v2.2.2^40,41^ to select unrelated individuals separated by at least two degrees of relatedness. After filtering, we have 1626 unrelated individuals in the FHS data and 2046 unrelated individuals in the JHS data. The purpose of removing relatives is to improve computational efficiency. Accuracy is unchanged when relatives are included (data not shown).

When phasing haplotypes, detecting IBD segments, and gap-filling IBD segments, we used a 1cM/Mb recombination rate. The IBD segments for our method were obtained by applying Refined IBD^42^ (LOD threshold = 1, minimum length 300kb) with gap-filling (maximum gap distance = 500kb, maximum number of discordant sites = 1). The thresholds (LOD 1 and minimum length 300kb) used in Refined IBD are quite low. However, in conjunction with the gap-fill step they allow the procedure to find as much IBD as possible, some of which will have a large genetic length and hence pass the subsequent filtering for IBD coverage (see the next section). The low thresholds used with Refined IBD will result in some short reported IBD segments that are actually the conflation of several shorter IBD segments^43^. However, for the purpose of estimating the genetic map, the benefit of the increased number of IBD segments is greater than the additional noise due to some IBD segment conflation. Use of a larger minimum physical length for IBD segments results in loss of accuracy (Figure S5).

The estimated recombination maps are normalized by the genetic length of each chromosome from the deCODE map, or by the true total genetic length for the simulated data. For comparison with our maps, we lifted over the AA map and the AfAdm map from build 36 to build 37 and the deCODE map from build 38 to build 37 using the following strategy. First we converted the target map to the bed interval and rate format, as “chr#:from-to rate”. Then we lifted over using the UCSC online tool: https://genome.ucsc.edu/cgi-bin/hgLiftOver, outputting the interval positions in bed format. We removed intervals that failed to be converted or for which the interval length changed by more than 1%. In total 133.7Mb was removed from the deCODE map, 139.6Mb was removed from the AA map, and 283.8Mb was removed from AfAdm map. Finally we mapped the recombination rates from each original map to the remaining intervals in build 37.

### Counting IBD ends to estimate recombination rates

One major issue in using IBD segments to estimate recombination rates is that IBD segments of a given physical length are not evenly distributed along the target genome when the recombination rates vary. To deal with this issue, we use a coverage threshold to make sure that the IBD endpoints for each interval use IBD segments drawn from equivalent distributions.

The IBD coverage of an interval (a genomic region of specified physical length) is the number of IBD segments covering the interval. IBD segments that partially cover the interval contribute a fractional value to the coverage equal to the proportion of the interval covered. The coverage is calculated for each interval, and the minimal value is determined. Then, in each interval, the segments with shortest genetic length are removed until removing an additional segment from the interval would reduce the coverage below this minimal level. An IBD segment may be removed from one interval but retained in another.

We use a constant recombination rate (1cM/Mb) to initiate the iterative estimation procedure. In each iteration, we re-estimate the genetic lengths of the IBD segments using the current recombination map, and we re-apply the coverage threshold. We then update the recombination rates for each interval based on the number of IBD endpoints located in the interval:

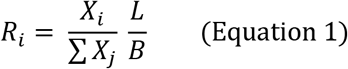

For the *i*-th interval, *R*_*i*_ is the estimated recombination rate, *X*_*i*_ is the number of IBD segment endpoints in the interval, *L* is the genetic length of the chromosome, and *B* is the physical length of the interval (the same for each interval). The genetic length of the chromosome is obtained from an external source such as a family-based genetic map.

In order to improve convergence, we use the average of the two previous estimates as the input recombination map to the next iteration (starting with the third iteration). Without this averaging, we found that the correlation between the true and simulated map oscillated up and down between successive iterations (Figure S6).

### Estimation at chromosome ends

We need to treat the ends of the chromosome differently, because IBD segments cannot continue beyond the end of the chromosome. Thus IBD segments starting or ending at a chromosome end are shorter on average, and fewer of these IBD segments will be detected. This results in a lack of right ends of IBD segments in intervals near the left end of the chromosome, and of left ends of IBD segments near the right end of the chromosome.

When estimating the recombination rate in an interval near the chromosome end, we make several changes to the algorithm described above. In order to describe these changes, we define chromosome end regions, and their neighboring adjunct regions (Figure 6). The end region starts at the chromosome end and has genetic length equal to the median genetic length of all IBD segments that extend to that chromosome end, plus any additional length required in order to have the end of this region correspond to a breakpoint between intervals. The remaining region between the two ending regions is the mid region. The adjunct region corresponding to an end region immediately follows the end region (on the side towards the middle of the chromosome) and has the same physical length as the end-region. During this procedure, we are not estimating the recombination rates of the intervals in the adjunct region. Rates in this region are estimated using the unmodified procedure described earlier. In what follows, we describe the changes to the algorithm with respect to the left end of the chromosome; the right end is analogous.

**Figure 6:**
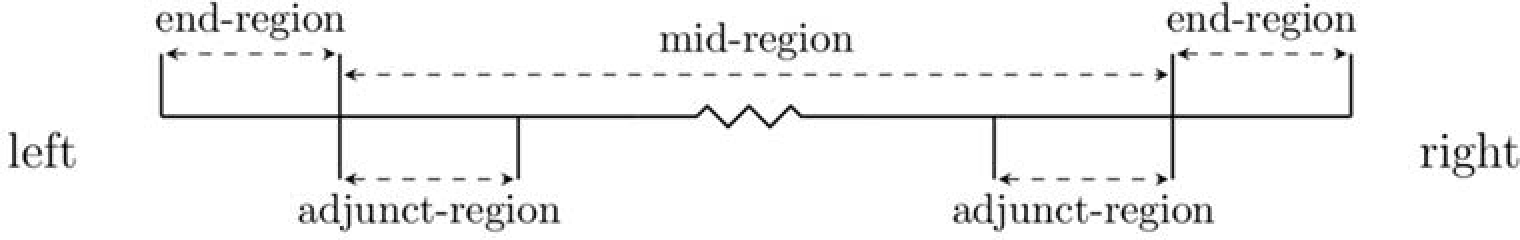
Chromosome regions for recombination rate estimation. The end region has genetic length equal to the median genetic length of IBD segments that extend to the chromosome end, and the adjunct-region is next to the end region and has the same genetic length as the end region.

The first change is that we count only the left ends of the IBD segments, rather than both endpoints of the IBD segments. This is because there will be a relative lack of right ends of IBD segments near the left chromosome end because many IBD segments that are censored by the left chromosome end will not be detected. In contrast, there will be no reduction in left ends of IBD segments close to the end of the chromosome.

The second change is that we need to modify the application of the IBD coverage threshold so that it has equal effect in all intervals in the end region, regardless of how close they are to the chromosome end. The left chromosome end left-censors the IBD segments that reach that chromosome end, so the visible lengths of the segments are shorter than they would otherwise be. For intervals other than the left-most one, we can mimic this censoring by removing those parts of IBD segments that fall beyond the left boundary of the interval. This trimming reduces the lengths of the IBD segments, and is performed only with respect to a given interval. The part of an IBD segment that is trimmed off when calculating segment lengths for one interval may be retained when calculating lengths for another interval. Thus, for each interval, not only for the left-most interval, the IBD segments that intersect the interval are left-censored by the left side of the interval. These adjusted IBD lengths are used when excluding the shortest IBD segments to equalize IBD coverage in each interval.

Recombination rates calculated with our method are relative. We use a user-specified total chromosome genetic length to normalize them. Since the estimation procedure for the end region and the mid region differ, we must put the two sets of results on the same scale. We do this by applying the end-region procedure for censoring IBD segments and equalizing IBD segment coverage to the adjunct region. Since we also have IBD end counts from the mid-region procedure for the adjunct region, we normalize the results from IBD end point counts for the end region by multiplying by the ratio of the IBD end counts in the adjunct region obtained from the mid-region and end-region methods.

Thus for intervals in the end region we obtain an estimate of what the two-sided end count would be if the interval was not affected by the chromosome end censoring:

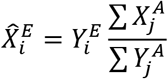

where 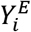 is the left-sided IBD end count for interval *i* in the left end-region; 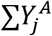 is the total count of left-sided IBD ends for intervals in the adjunct region, applying the end-region algorithm described in this section; and 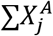 is the total count of IBD ends (left and right) for intervals in the adjunct region, based on the original (mid-region) algorithm. The value 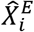 is the adjusted end count for interval *i* in the end-region; this is used in place of *X*_*i*_ in the recombination rate estimation formula (Equation 1).

### Fine-scale estimation

We have proposed a procedure for estimating recombination rates from IBD endpoints in the previous two sections. This procedure works well when the number of detected IBD segments is large due to a large sample size, and the interval size is large. However, when the interval size decreases for fine-scale estimation, the coverage threshold tends to decrease, resulting in loss of information at small sample sizes. We thus improve fine scale estimation by running our algorithm in two steps. First, we construct a recombination map at a large scale, for example with an interval size of 500kb (which we use as the at the first step scale in all the analyses in this paper). We obtain estimates of genetic length for each large interval, and we fix these large-scale lengths in the second step. In the second step, we divide each large interval into many smaller sub-intervals at the desired scale. For example, if results at a 10 kb scale are desired, sub-intervals of length 10 kb are used. The estimation procedure for these short intervals is slightly modified from the algorithm described above.

For the fine-scale estimation, the IBD coverage threshold is based on the minimal coverage of the sub-intervals within a large interval, rather than on the minimal coverage of intervals across the whole chromosome. The local coverage threshold tends to be larger than the global threshold used in the large-scale estimation because there is typically less variability in recombination rate across an interval than across a whole chromosome.

After the large-scale estimation, the lengths of the large intervals in the end region are known and it is no longer necessary to use an adjunct region to normalize lengths in the end region. However, we do still need to use only the one-sided IBD end counts, and to censor the IBD segments intersecting each interval when applying the IBD coverage threshold in the end region. As in the large-scale step, we adjust the genetic lengths by trimming off that part of the IBD segment that extends beyond the sub-interval in the direction of the nearby chromosome end.

Within each large interval (whether within an end region or not), we estimate the recombination rates of the sub-intervals using the formula in Equation 1, using the previously calculated genetic length of the large interval as the region length *L*. For intervals that are not in the end regions, we use two-sided IBD end counts for the *X_i_*, while for intervals within the two end regions, we replace these with the one-sided IBD end counts.

We have implemented this two-step procedure in the IBDrecomb program, and the fine scale estimation step is automatically applied when the fine interval size parameter (for the second stage of estimation) is set to a value that is smaller than the large interval size parameter (for the first stage of estimation; 500kb by default).

## Online Resources

IBDrecomb (including FHS and JHS maps): https://github.com/YingZhou001/IBDrecomb

Msprime: https://msprime.readthedocs.io/en/stable

Refined IBD and Gap-filling tool: http://faculty.washington.edu/browning/refined-ibd.html

AA map (build 36): http://www.well.ox.ac.uk/~anjali/AAmap/aamap.tar.gz

AfAdm map (build 36): https://www.eeb.ucla.edu/Faculty/Novembre/software/AfricanAmerican_AfricanCaribbean_recombination_maps.zip

deCODE map (build 38): https://science.sciencemag.org/highwire/filestream/721792/field_highwire_adjunct_files/4/aau1043_DataS3.gz

Hapmap II combined map (build 37): ftp://ftp.ncbi.nlm.nih.gov/hapmap/recombination/2011-01_phaseII_B37/genetic_map_HapMapII_GRCh37.tar.gz

1000 genome maps (build 37): ftp://ftp.1000genomes.ebi.ac.uk/vol1/ftp/technical/working/20130507_omni_recombination_rates/

Bhérer et al’s refined maps (build 37): https://github.com/cbherer/Bherer_etal_SexualDimorphismRecombination/raw/master

## ACKNOWLEDGMENTS

This work was supported by NIH grant R01HG005701. Whole genome sequencing (WGS) for the Trans-Omics in Precision Medicine (TOPMed) program was supported by the National Heart, Lung and Blood Institute (NHLBI). WGS for “NHLBI TOPMed: Whole Genome Sequencing and Related Phenotypes in the Framingham Heart Study” (phs000974.v2.p2) was performed at the Broad Institute of MIT and Harvard (HHSN268201500014C). WGS for “NHLBI TOPMed: The Jackson Heart Study” (phs000964.v2.p1) was performed at the University of Washington Northwest Genomics Center (HHSN268201100037C). Centralized read mapping and genotype calling, along with variant quality metrics and filtering were provided by the TOPMed Informatics Research Center (3R01HL-117626-02S1). Phenotype harmonization, data management, sample-identity QC, and general study coordination, were provided by the TOPMed Data Coordinating Center (3R01HL-120393-02S1). We gratefully acknowledge the studies and participants who provided biological samples and data for TOPMed. The Jackson Heart Study (JHS) is supported and conducted in collaboration with Jackson State University (HHSN268201800013I), Tougaloo College (HHSN268201800014I), the Mississippi State Department of Health (HHSN268201800015I/HHSN26800001) and the University of Mississippi Medical Center (HHSN268201800010I, HHSN268201800011I and HHSN268201800012I) contracts from the National Heart, Lung, and Blood Institute (NHLBI) and the National Institute for Minority Health and Health Disparities (NIMHD). The Framingham Heart Study is conducted and supported by the National Heart, Lung, and Blood Institute (NHLBI) in collaboration with Boston University (Contract No. N01-HC-25195 and HHSN268201500001I). This manuscript was not prepared in collaboration with investigators of the Framingham Heart Study or the Jackson Heart Study and does not necessarily reflect the opinions or views of these studies or of the NHLBI.

## AUTHOR CONTRIBUTIONS

Y.Z. conceived the study, developed methodology and software, performed analyses, and co-wrote the paper. B.L.B. provided input into the methodology and edited the paper. S.R.B. supervised the study, provided input into the methodology, and co-wrote the paper.

